# Pre-saccadic Preview Shapes Post-Saccadic Processing More Where Perception is Poor

**DOI:** 10.1101/2023.05.18.541028

**Authors:** Xiaoyi Liu, David Melcher, Marisa Carrasco, Nina M. Hanning

## Abstract

The pre-saccadic preview of a peripheral target enhances the efficiency of its post-saccadic processing, termed the extrafoveal preview effect. Peripheral visual performance –and thus the quality of the preview– varies around the visual field, even at iso-eccentric locations: it is better along the horizontal than vertical meridian and along the lower than upper vertical meridian. To investigate whether these polar angle asymmetries influence the preview effect, we asked human participants (to preview four tilted gratings at the cardinals, until a central cue indicated to which one to saccade. During the saccade, the target orientation either remained or slightly changed (valid/invalid preview). After saccade landing, participants discriminated the orientation of the (briefly presented) second grating. Stimulus contrast was titrated with adaptive staircases to assess visual performance. Expectedly, valid previews increased participants’ post-saccadic contrast sensitivity. This preview benefit, however, was inversely related to polar angle perceptual asymmetries; largest at the upper, and smallest at the horizontal meridian. This finding reveals that the visual system compensates for peripheral asymmetries when integrating information across saccades, by selectively assigning higher weights to the less-well perceived preview information. Our study supports the recent line of evidence showing that perceptual dynamics around saccades vary with eye movement direction.

**Significance Statement:** We constantly make saccadic eye movements to bring relevant visual information into the fovea, which has the highest acuity. Before each saccade, we use “previewed” peripheral information to support our post-saccadic vision. Our sensitivity varies around the visual field –at the same eccentricity it is best along the horizontal meridian and worst at the upper vertical meridian. An optimal visual system should rely more on previewed information with higher precision. Our study reveals the opposite: peripheral preview shapes subsequent post-saccadic foveal processing more at locations where peripheral vision is worse. This finding implies that the human visual system compensates for sensitivity differences around the visual field when integrating information across eye movements.

## Introduction

When exploring their surroundings, humans make fast saccadic eye movements to bring different parts of the scene onto the fovea, the central region of the retina with the highest visual acuity. Despite its low resolution, the *preview* of peripheral visual information plays a critical role in preparing and guiding saccades (1, 2). Inputs previewed *before* the saccade are integrated with the *post*-saccadic percepts, facilitating subsequent foveal processing (review: 3). This *extrafoveal preview effect* occurs with a variety of stimuli, including low-level features (4, 5), words (6), and objects (7–9). It is reflected both at the behavioral level –e.g., higher accuracy and faster reaction times (e.g., 7) – and at the neural level –e.g., faster decoding time (10).

The preview effect has been studied exclusively along the horizontal meridian (HM), but peripheral vision varies greatly with polar angle: it is better along the horizontal than vertical meridian (HM>VM: horizontal-vertical anisotropy; HVA) and along the lower than upper vertical meridian (LVM>UVM: vertical meridian asymmetry; VMA) (11–13). Visual sensitivity can be two to three times higher for stimuli on the HM than VM (14, 15). These polar angle asymmetries are present in many visual tasks, such as contrast sensitivity (12, 14, 16, 17), visual acuity (18, 19), spatial resolution (20, 21), and crowding (13, 22) (For a review on polar angle asymmetries, see 23). However, it is unknown whether and how our heterogeneous peripheral vision affects the integration of pre- and post-saccadic information for different saccade directions.

Here we consider two opposite hypotheses. According to the optimal integration framework (24–26), trans-saccadic integration is similar to other types of sensory integration across modalities, during which the brain integrates information by weighing the sensory evidence based on relative precision. Estimated reliability is maximized when stronger sensory signals (e.g., higher signal-to-noise ratio, reduced uncertainty) receive higher weights (25). Correspondingly, preview benefits in reading are reduced or extinguished when the preview word is located at a farther eccentricity, e.g., two versus one word from the current fixation (27, 28). More evidence comes from gaze-contingent tasks that present stimuli according to the observers’ gaze position and systematically vary the relative visibility of peripheral and foveated targets by changing either the stimulus contrast (29) or eccentricity (9), or by adding noise (9, 30). These studies have revealed that with incongruent pre- and post-saccadic percepts, trans-saccadic perceptual judgements are increasingly biased towards the peripheral preview as its visibility increases. Because object visibility also varies with polar angle, one could expect the magnitude of the preview effect around the visual field to follow the same pattern: a larger preview effect on the HM, where peripheral vision is best, followed by the lower- and then the upper-vertical meridian.

In contrast, the visual system might compensate for polar angle asymmetries in peripheral vision when integrating information across saccades, enhancing post-saccadic vision more at locations where it is needed the most. Endogenous attention, the voluntary allocation of processing resources based on one’s goals (31), is an example of this process. Depending on the current task demand, endogenous attention can exert different perceptual effects (e.g., 32, 33). For example, it either increases or decreases the temporal integration window when observers are instructed to segregate or integrate two closely presented stimuli. This effect is more pronounced when baseline performance is worse (32). A similar compensation mechanism during trans-saccadic integration would give higher weights to the pre-saccadic preview signals at locations where peripheral vision is worse, leading to equal preview effects around the visual field, or even a reversed pattern of preview asymmetries.

However, polar angle asymmetries in peripheral vision are resilient; they are not alleviated by covert temporal (34) or spatial attention: Both exogenous attention –which is involuntary, fast and transient (12, 35)– and endogenous attention –which is voluntary, slower and sustained (36)– similarly improve performance around the visual field, thereby preserving the polar angle asymmetries. Pre-saccadic attention –which is automatically deployed to the target of an upcoming eye movement during saccade preparation (37–43) – enhances perception similarly around the visual field (19) or even exacerbates polar angle asymmetries, by benefitting performance less where it is already worse (44, 45). Thus, neither covert nor pre-saccadic attention overcome polar angle asymmetries in peripheral vision.

Pre-saccadic attention contributes to trans-saccadic integration, but the latter entails more processes (3, 7) and relies on a prolonged interaction among the oculomotor network and the visual system before and after saccades (46, 47). Therefore, to investigate whether and how the extrafoveal preview effect, a measure for trans-saccadic integration, interacts with polar angle asymmetries in peripheral vision, we first assessed participants’ baseline contrast sensitivity at four isoeccentric cardinal locations during fixation, and then measured their preview effect at the same locations with an *integration* task, as the difference in logarithmic post-saccadic contrast sensitivity following valid vs. invalid previews.

Our findings are consistent with the compensation hypothesis. Unlike previous reports that polar angle asymmetries are preserved or even exacerbated by covert and pre-saccadic attention (e.g., 12, 19, 36, 42, 43), we found the largest preview effect at the UVM, followed by the LVM, and lastly the HM –inversely correlated with peripheral polar angle asymmetries during fixation.

## Results

### Baseline sensitivity during fixation varies with polar angle

To establish the typical polar angle asymmetries (horizontal-vertical anisotropy, HVA; and vertical meridian asymmetry, VMA), participants first completed a baseline *fixation* task, in which four gratings (one tilted target with three vertical distractors) were briefly presented 8 dva (degree of visual angle) above, below, left, and right of the central fixation (**Fig. 1a**). Participants indicated the tilt direction of the target. Stimulus contrast was adjusted trial-by-trial via an adaptive staircase procedure (PEST; see Materials and Methods for details).

**Figure 1.**
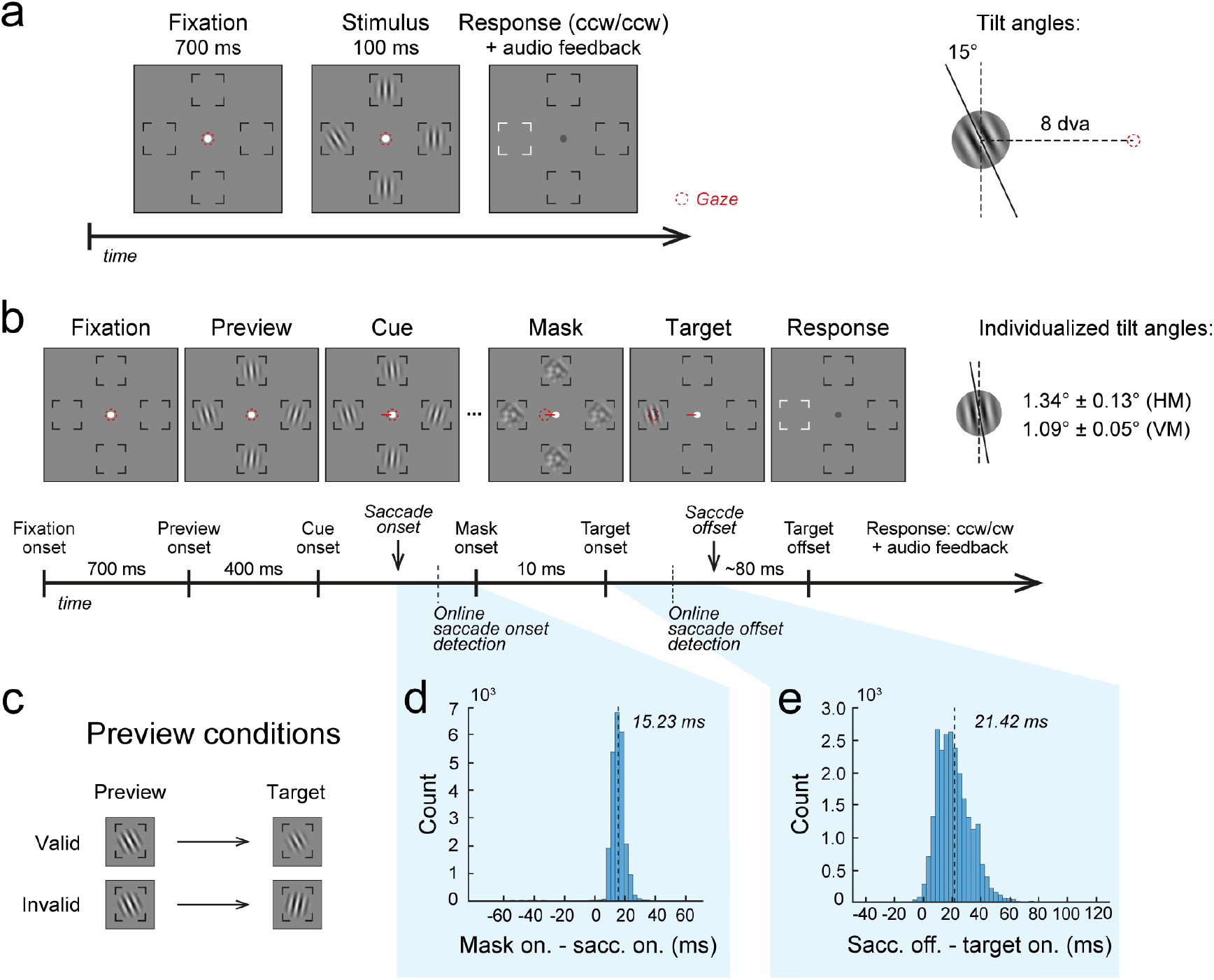
Illustration of the task. **a**. *Fixation* task. Each trial started with a central fixation dot, followed by the stimuli. Participants indicated the tilt direction of a tilted Gabor (±15° relative to vertical) presented among distractors. **b**. Integration task. After fixation, participants previewed four slightly tilted Gabors, followed by a saccade cue for an immediate eye movement. Saccade onset, detected by an online algorithm (see Methods), triggered a 10ms mask, followed by the target. The target disappeared ∼80ms after saccade offset. Participants then indicated the tilt direction of the second, foveated target. The tilt angles of Gabor stimuli were determined by a titration task (see SI) **c**. Task conditions in the integration task. **d**. Temporal delay between offline detected saccade onset and mask onset. **e**. Temporal delay between target onset and offline detected saccade offset.

Consistent with the HVA, a paired-sample t-test revealed a significant difference in contrast sensitivity between the horizontal and vertical meridian (H01: horizontal = vertical), *t*(13) = 3.40, *p* = .0047, BF = 10.51, Cohen’s d = 0.91, CI_*d*_ = [0.33, 1.49]. Sensitivity along the HM (33.13 ± 4.08) (M ± SE) was significantly higher than that along the VM (21.73 ± 2.39). A Repeated-measures ANOVA with location (left, right, upper, lower) as within-subject factor (H02: left = right = upper = lower) showed a main effect of location, *F* (3, 39) = 8.94, *p* < .001, BF > 100, *η*_*p*_^2^ = .41, 95% CI_*η*_ = [.14, .55]. Consistent with the VMA, post-hoc tests showed no difference between the left (32.47 ± 4.09) and right (33.79 ± 4.4) HM (BF = .31), whereas sensitivity at the lower VM (24.96 ± 3.1) was significantly higher than the upper VM (18.49 ± 2.19), *t*(13) = 2.64, *p* = .041, BF = 3.19, Cohen’s d = 0.71, CI_*d*_ = [0.13, 1.28].

To visualize the individual differences in HVA and VMA, we plotted each participant’s contrast sensitivity along the HM against the VM (**Fig. 2b**), and along the LVM against the UVM (**Fig. 2c**). Diagonal lines indicate equivalent performance. HVA and VMA were found in all participants, except in two for HVA and four for VMA. The strength of the asymmetries was computed as the sensitivity difference between the two respective meridians divided by their average, with a positive number representing a typical HVA/VMA. The correlation between HVA and VMA was not significant (*r* = .20, *p* = .50), suggesting (at least partially) different origins (14, 18).

**Figure 2.**
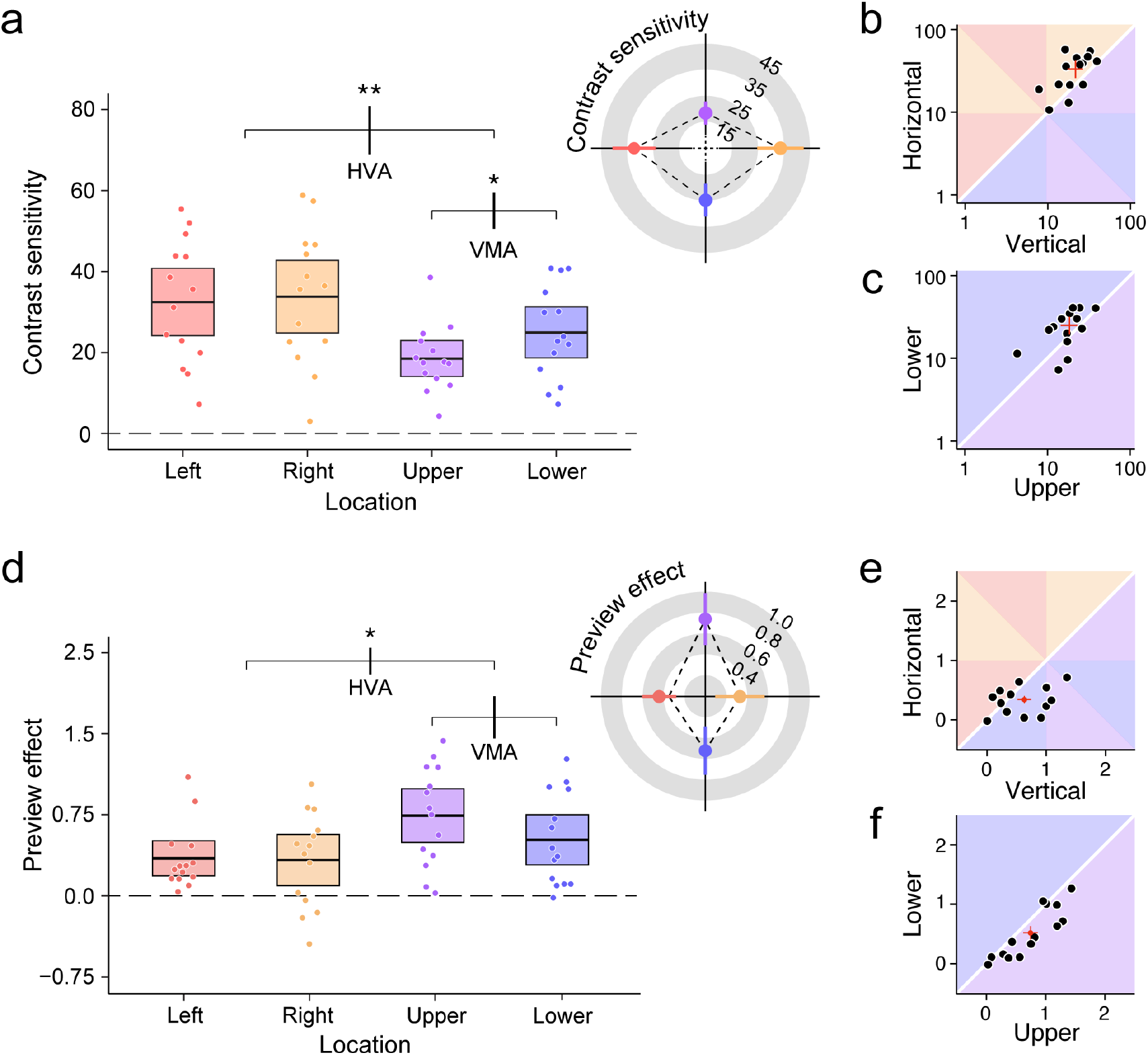
Behavioral results in the *fixation* and *integration* tasks. **a.d**. Contrast sensitivity in the *fixation* task computed as the reciprocal of the staircase threshold estimate (**a**) or preview effect in the *integration* task computed as the difference in log-sensitivity between the valid and invalid conditions (**d**) as a function of target location. Boxes depict the 95% CIs around the mean, represented by the black lines. Horizontal brackets indicate comparisons between the horizontal vs. vertical meridian (HVA) and upper vs. lower vertical meridian (VMA); error bars on the brackets represent 95% CIs of the difference between compared conditions. * *p* < .05, ** *p* < .01 (after correcting for multiple comparisons). Polar plots depict the same data. **b.e**. Individual participants’ contrast sensitivity (**b**) or preview effect (**e**) along the horizontal meridian plotted against the vertical meridian. Red error bars represent 95% Cis of group averages. **c.f**. Individual participants’ contrast sensitivity (**c**) or preview effect (**f**) along the lower against the upper vertical meridian.

### The preview effect is inversely correlated with peripheral sensitivity

The same participants then performed an *integration* task: They peripherally viewed four tilted gratings before a saccade cue indicated to which grating an immediate saccade had to be made. During the saccade, the grating at the saccade target either remained identical (*valid* condition), or flipped its tilt angle (e.g., from +1° to -1°) (*invalid* condition; **Fig. 1d**). After the saccade, participants indicated the tilt orientation (clockwise vs. counterclockwise) of the post-saccadic target grating (see Materials and Methods for details).

We quantified the magnitude of the preview effect as the difference in logarithmic sensitivity between the valid and invalid conditions. Strikingly, the HVA and VMA in the preview effect showed the opposite pattern to peripheral sensitivity in the baseline task: A paired-sample t-test showed a significant effect of meridian, *t*(13) = -2.67, *p* = .019, BF = 3.35, Cohen’s d = 0.71, CI_*d*_ = [0.14, 1.29], where the preview effect was *larger* at the VM (0.63 ± 0.08) than the HM (0.34 ± 0.07), (**Fig. 2d;** see **Fig. S1** for sensitivity in the valid and invalid conditions separately) – i.e., inversely related to polar angle differences in peripheral vision (HVA, *baseline* task). A repeated-measures ANOVA with location as within-subject factor revealed a main effect of location, *F* (3, 39) = 4.28, *p* = .011, BF = 6.17, *η*_*p*_^2^ = .25, 95% CI_*η*_ = [.02, .41]. Post-hoc tests showed no difference between the left (0.35 ± 0.08) and right (0.33 ± 0.12) HM (BF = .27). The preview effect at the upper VM (0.74 ± 0.12), however, was significantly *smaller* than at the lower VM (0.52 ± 0.11), *t*(13) = 3.79, *p* = .005, BF = 19.63, Cohen’s d = 1.01, CI_*d*_ = [0.44, 1.59] – likewise showing the inverse pattern as polar angle differences in peripheral vision (VMA, *baseline* task). Both the reverse VMA in the preview effect were evident at the individual observer level (**Fig. 2e,f**)).

To further evaluate the relation between the peripheral contrast sensitivity during fixation and the magnitude of the preview effect at the individual level, we conducted a repeated-measures correlation (48), which accounts for the violation of independence in data due to repeated within-participant data points. This way we measured the variability across target locations by removing the inter-individual variability. Aligned with the reversed pattern of asymmetries indicated by the ANOVA, we found a significant negative intra-individual correlation between peripheral contrast sensitivity and the preview effect, *r*_*m*_ = -.41, *p* = .007, BF = 24.58, 95% CI = [-0.63, -.12] (**Fig. 3**). Thus, within individuals, the preview effect was larger where their peripheral sensitivity was lower (e.g., UVM).

**Figure 3.**
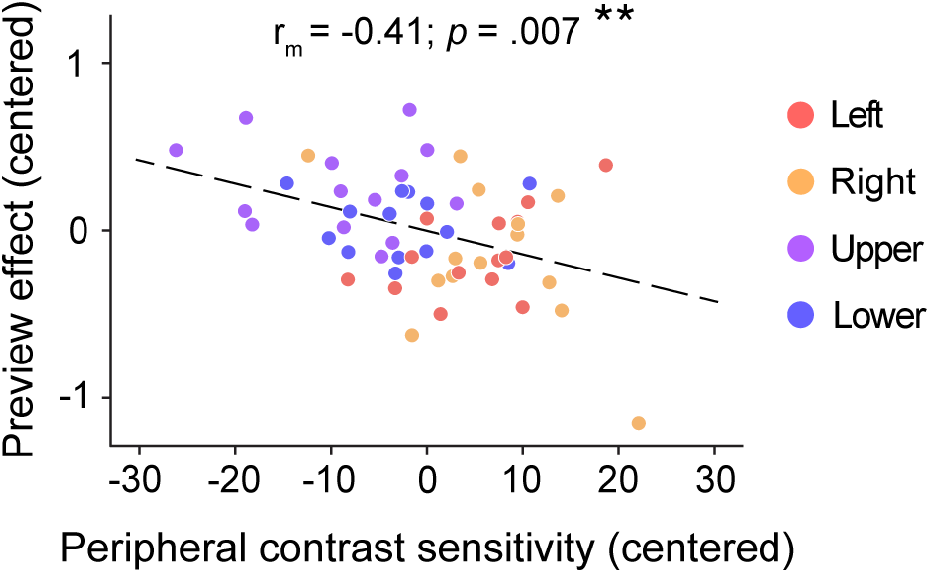
Individual participants’ preview effect as a function of peripheral contrast sensitivity. Data centered to each participants’ average across target locations. Dots represent individual data at the four cardinal locations. The regression line represents the intra-individual (repeated-measures) correlation.

### Lowest post-saccadic sensitivity at the UVM

The same participants also participated in a *postview* task, which measured pure post-saccadic foveal contrast sensitivity, without pre-saccadic preview (the procedure otherwise matched the *integration* task, see Materials and Methods for details). We expected equivalent performance following saccades in different directions, as the briefly presented target was always foveated and polar angle asymmetries therefore should not affect perception. Surprisingly, we found a main effect of location, *F* (3, 39) = 4.34, *p* = .010, *η*_*p*_^2^ = .25, 95% CI_*η*_ = [.02, .41], which was driven by a lower post-saccadic contrast sensitivity *after* upward saccades (15.00 ± 3.16) compared to the other directions (rightward: 29.49 ± 4.76; *p* = .015, 95% CI = [0.91, 28.05]; leftward: 26.23 ± 3.64; *p* = .054, 95% CI = [-2.34, 24.80], marginally significant; downward: 30.85 ± 2.77; *p* = .014, 95% CI = [2.28, 29.42]) (**Fig. 4**). Other post-hoc contrasts did not reach significance.

**Figure 4.**
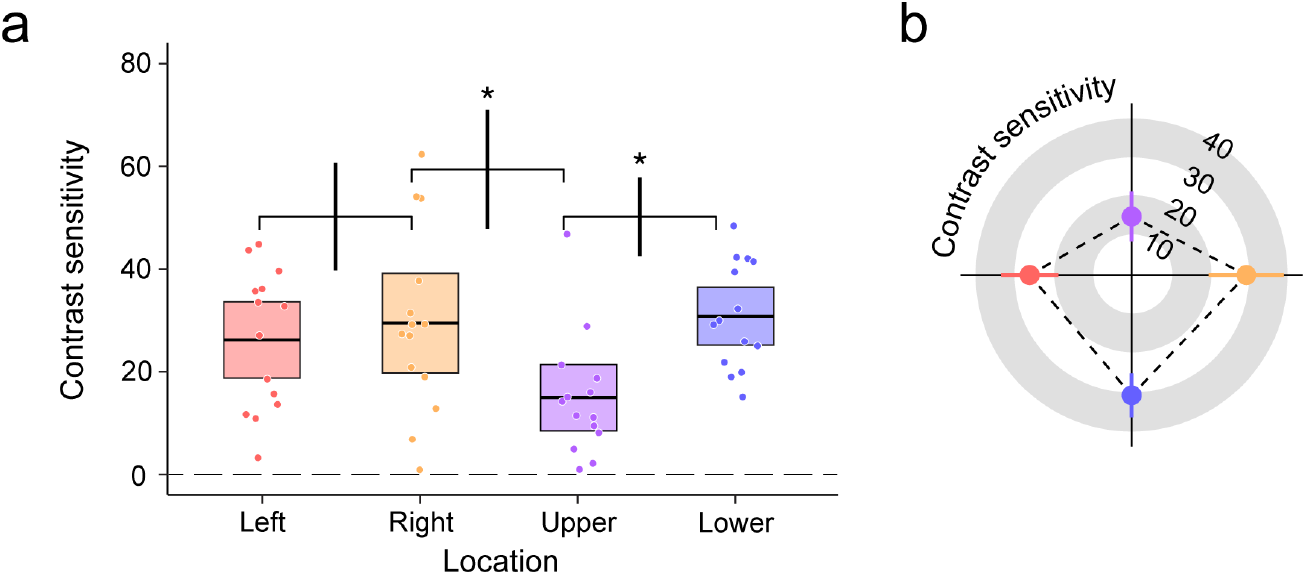
Contrast sensitivity in the *postview* task as a function of target location. **a**. Boxes depicts the 95% CIs around the mean, represented by black lines. Horizontal brackets indicate comparisons between the respective target location, with the error bars on the brackets representing 95% CIs of the difference between the compared conditions. * *p* < .05 (after correcting for multiple comparisons). **b**. Polar plots showing the same data.

Importantly, post-saccadic sensitivity was not predictive of the preview effect: A linear regression model (M1) with peripheral contrast sensitivity and post-saccadic foveal sensitivity as predictors for the preview effect showed that peripheral sensitivity (*baseline* task) predicted the preview effect in a negative way (*p* = .03; consistent with the repeated-measures correlation, **Fig. 3**), but post-saccadic sensitivity (*postview* task), was not associated with the preview effect. We ran a second regression model with peripheral sensitivity as the sole predictor (M2). The comparison of M1 and M2 via a likelihood ratio test showed that peripheral and post-saccadic sensitivity combined (M1) did not explain more variance in the preview effect than peripheral sensitivity alone (M2; *p* = .15).

### Saccade parameters cannot explain the preview effect asymmetries

We verified that differences in saccade parameters (i.e., saccade latency, amplitude, precision, post-saccadic oscillations) did not give rise to the observed preview asymmetries around the visual field. Longer saccade latencies (and thus a longer preview duration) as well as less precise saccades or higher post-saccadic movement rates (and thus a lower quality postview) could lead participants to rely more on the preview information, thereby causing a larger preview effect. We therefore compared the eye movement data (collected in the *integration* and *postview* task) with 4 (saccade direction: left, right, upper, lower) × 3 (viewing condition: valid preview, invalid preview, postview) repeated-measures ANOVAs.

We found main effects of saccade direction, *F* (3, 39) = 32.14, *p* < .001, *η*_*p*_^2^ = .71, 95% CI_*η*_ = [.51, .79] and viewing condition, *F* (2, 26) = 15.94, *p* < .001, *η*_*p*_^2^ = .55, 95% CI_*η*_ = [.24, .69] (**Fig. 5a**) on saccade latency. Latencies were significantly longer for downward saccades (215.05 ± 2.6 ms) than saccades in the other directions (left: 194.91 ± 1.98, 95% CI = [13.60, 26.69]; right: 197.81 ± 1.88, 95% CI = [10.64, 23.73]; upper: 196.22 ± 2.13 ms, 95% CI = [12.29, 25.38]; all *p* < .001), consistent with previous studies (19, 44, 45, 49). Moreover, latencies were shorter in the invalid (195.85 ± 1.87 ms) than in the valid preview (203.99 ± 2.21 ms; *p* < .001, 95% CI = [-12.21, - 4.07]) and postview (203.20 ± 2.24 ms; *p* < .001, 95% CI = [-11.42, -3.28]) viewing conditions. As the perceptual task was relatively harder for the invalid preview condition, Gabor contrast (adjusted by the staircase) accordingly was higher (invalid preview: 21.24% ± 3.42%; valid preview: 4.58% ±0.56%; postview: 9.49% ±2.45%), which might explain this latency difference. Neither saccade amplitudes (7.86 ± 0.03 dva; **Fig. 5b**) nor saccade precision (i.e., the Euclidean distance between the target and saccade endpoint; 0.97 ± 0.01 dva; **Fig. 5c**) differed as a function of target location or viewing condition.

**Figure 5.**
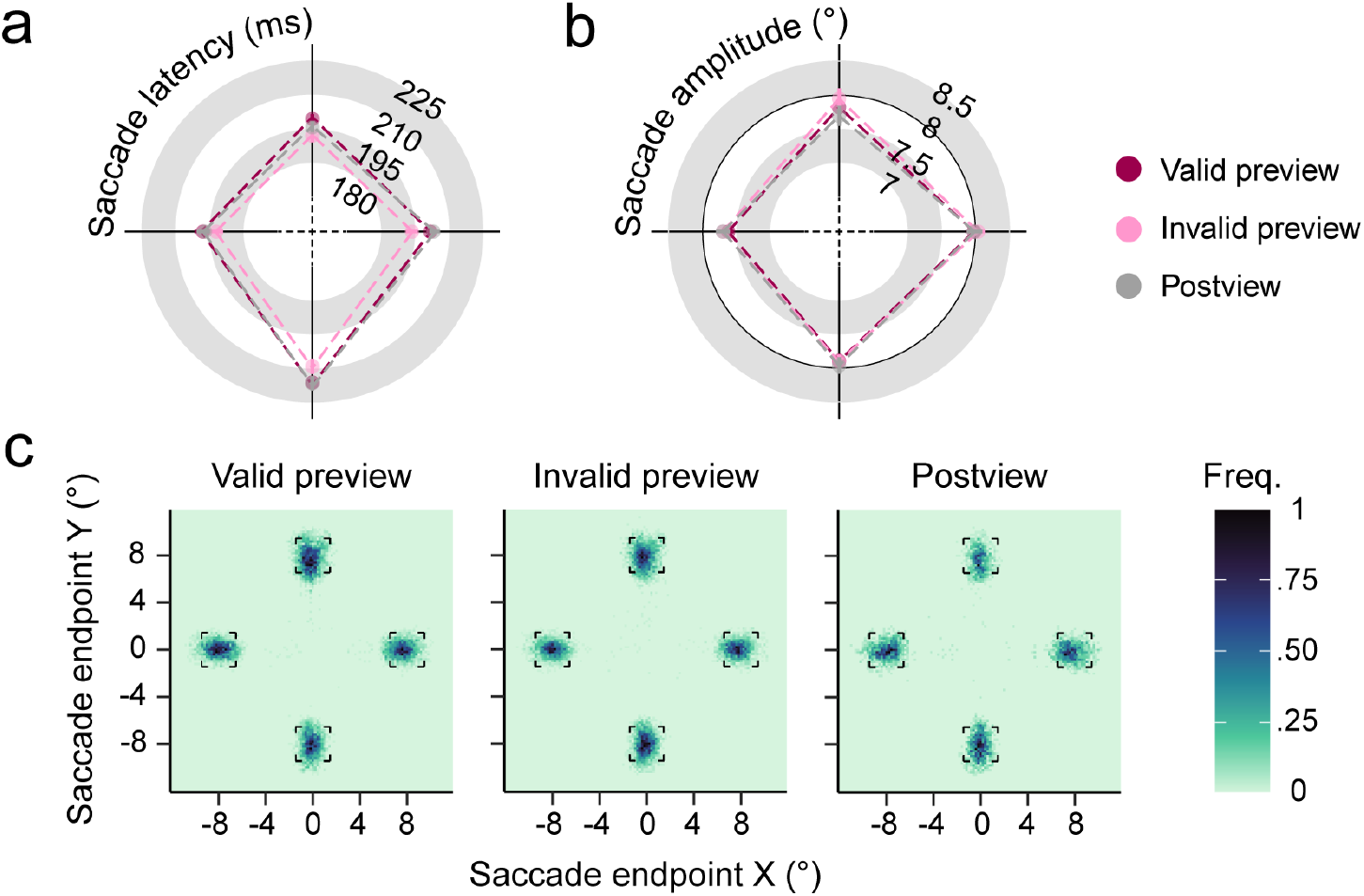
Group averaged saccade latency (**a**) and saccade amplitude (**b**) as a function of saccade direction and target location. Error bars indicate 95% CIs. **c**. Saccade endpoint frequency maps averaged across participants depicting saccade landing error. The outlined squares represent the placeholders.

#### Post-saccadic eye movements

Shortly after a saccade, gaze is unstable. The eyes may drift or participants may perform tiny (corrective) saccades. As any post-saccadic movement shifts the image on the retina and thereby directly affects foveal perception (50), we evaluated participants’ post-saccadic eye movements and its potential effect on the preview effect. To consider any ocular movements after saccade landing, including post-saccadic oscillations (i.e., ocular instability due to pupil movements; 49), fixational eye movements (micro-saccades) and ocular drift, we analyzed eye movement rate and distance gaze covered (see Supporting Information SI) during the post-saccadic target presentation with repeated-measures ANOVAs. Post-saccadic movement distance did not differ between saccade directions or viewing conditions, with gaze samples covering an average range of 0.51 dva (± 0.03 dva) (**Fig. 6a**).

**Figure 6.**
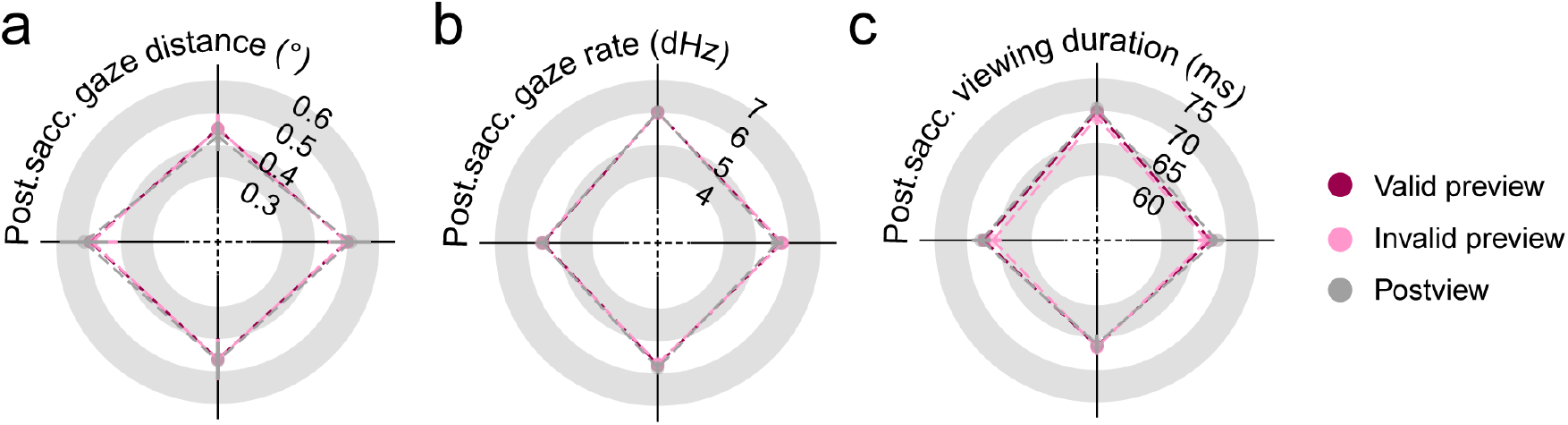
Group averaged post-saccadic gaze distance (**a**), saccade rate (**b**), and target viewing duration (**c**) as a function of task condition and saccade direction. Error bars indicate 95% CIs.

There was a main effect of saccade direction on post-saccadic eye movement rate, *F* (3, 39) = 3.11, *p* = .037, *η*_*p*_^2^ = .19, 95% CI_*η*_ = [.00, .35], driven by a higher movement rate after upward (6.02 ± 0.07 Hz) than leftward (5.56 ± 0.08 Hz, *p* = .025, 95% CI = [.040, .87]) saccades (**Fig. 6b**). Because other post-hoc contrasts did not reach significance, differences in post-saccadic ocular instability can only partially explain the observed asymmetries in preview effect and post-saccadic sensitivity.

#### Post-saccadic target viewing duration

Another potential source of the observed preview asymmetries could be the post-saccadic target viewing duration. A shorter viewing time could bias trans-saccadic integration toward the preview, leading to a larger preview effect. Even though we timed stimulus offset based on the estimated saccade offset (using an online criterion, see Methods), the actual target viewing time could have varied as a function of target location due to differences in saccade velocity or post-saccadic instability that could not be detected online. Indeed, there was a main effect of saccade direction, *F* (3, 39) = 4.31, *p* = .010, *η*_*p*_^2^ = .25, 95% CI_*η*_ = [.018, .41]. However, target viewing duration was slightly but significantly *longer* after upward saccades (69.78 ± 0.63ms) than leftward (67.18 ± 0.77ms; *p* = .039, 95% CI = [-.17, 5.37]) and downward (66.34 ± 0.73ms; *p* = .0082, 95% CI = [.67, 6.21]) saccades (**Fig. 6c**), ruling out the possibility that a *shorter* post-saccadic target duration time contributed to the larger preview effect at the UVM. There was also a main effect of viewing condition, *F* (2, 26) = 3.58, *p* = .042, *η*_*p*_^2^ = .22, 95% CI_*η*_ = [.00, .42], with a shorter target viewing found in the invalid preview (67.18 ± 0.64ms) than *postview* condition (68.25 ± 0.69ms; p = .041, 95% CI = [.033, 2.11])

## Discussion

We investigated trans-saccadic integration around the visual field, by testing whether and how the extrafoveal preview effect is modulated by polar angle asymmetries in peripheral vision. We replicated the typical polar angle asymmetries during fixation –higher contrast sensitivity at the horizontal than vertical meridian (HVA) and at the lower than upper vertical meridian (VMA) (11–13)– and found a robust preview effect at all cardinals: Post-saccadic foveal sensitivity was higher after a valid than invalid preview. Importantly, the magnitude of this preview effect was inversely related to the polar angle asymmetries: larger at locations with lower peripheral sensitivity. These preview asymmetries could not be explained by differences in saccade parameters. Our novel findings suggest that the visual system compensates for anticipated asymmetries in peripheral vision by assigning different weights to the previewed information across saccadic eye movements.

What could give rise to these preview asymmetries? Right before saccade onset, contrast sensitivity rapidly increases at the saccade target (41, 44, 45, 52) and renders the peripheral information more fovea-like already before the eyes start moving (19, 53–56). This pre-saccadic shift of attention allows a smooth handoff of pre-saccadic information to the post-saccadic processing area, which is necessary for continuity in trans-saccadic perception (57–61). However, previous studies have found an absent or reduced presaccadic benefit before upward saccades compared to saccades to other directions (44, 45). This could explain the worse post-saccadic foveal discrimination in the *postview* task after upward saccades than saccades in other directions: Without attention shifting prospectively to the saccade target –easing trans-saccadic integration by rendering the peripheral visual input more fovea-like– visual sensitivity might recover more slowly after saccade offset. Similarly, in the *integration* task, because the target disappeared briefly after saccade offset (before vision could fully recover), participants needed to rely more on the peripheral preview (62), leading to a larger preview effect at the UVM. Regardless of the mechanism underlying the worsened post-saccadic sensitivity following upward saccades, the visual system might have given more decision weight to the peripheral preview when a less reliable post-saccadic percepts was expected, consistent with the optimal integration framework (29, 60, 63). This differential weight assignment, however, cannot explain the reverse VMA in the preview effect, as the observed post-saccadic foveal sensitivity was equivalent between the LVM and HM. Our regression analysis further verified that it could not explain the variance in the preview effect, arguing against the optimal integration hypothesis.

Another potential mechanism that could have cause the observed preview asymmetries is pre-saccadic “foveal remapping”: Neuroimaging studies have found that before a saccade, neurons that process foveal information shift their receptive fields to also process detailed information about the peripheral saccade target (64, 65). Likewise, psychophysical evidence has shown that foveal processing of the saccade target begins already before saccade onset (66). Akin to presaccadic attention, this mechanism facilitates visual processing at the saccade target, and thereby a smooth transition from pre-to post-saccadic vision. Our visual system might strategically maximize foveal remapping for saccade targets with comparatively lower visual sensitivity (e.g., the LVM and UVM) to compensate for polar angle asymmetries in peripheral vision, leading to the observed reversed preview asymmetries.

Alternatively, the preview asymmetries might arise from the very first stage of trans-saccadic integration – when a prediction about the saccadic target is made based on peripheral inputs long before saccade onset (7, 58, 67). A stronger prediction about the peripheral saccade target could cause its (relative) up-weighting, leading to a larger preview effect. According to predictive processing, the preview effect is a measure of how much post-saccadic perception is biased toward a pre-saccadic prediction about the target. Although stronger prediction is intuitively associated with a higher peripheral sensitivity, strength and content of pre-saccadic predictions are learned over time: The repeated exposure to consistently invalid previews (e.g., an orange always changes to a basketball during a saccade), causes participants to make predictions about the saccadic target in the learnt direction, reducing the preview effect in response to statistical regularity (3, 68). Future research is needed to investigate how pre-saccadic prediction strength and content change around the visual field and how it is related to the preview asymmetries.

Our study is the first to investigate how heterogeneously perceived peripheral information is integrated across saccadic eye movements and differs from previous studies testing how object visibility affects trans-saccadic integration (9, 29) in an important way: Instead of external manipulation on stimulus attributes, the perceptual quality of peripheral stimuli in the current study was constrained by our visual system -an internal factor. On the one hand, these asymmetries in peripheral vision are so robust that they cannot be overridden by any attention mechanism studied so far (e.g., 12, 19, 34, 36, 42, 43). On the other hand, the current results indicate that our brain compensates for these asymmetries during trans-saccadic integration.

These results have significant implications in real-world applications. When driving, for example, people tend to fixate on one point while covertly monitoring the periphery. Polar angle asymmetries in peripheral vision suggest that drivers will be the most sensitive to objects along the HM (e.g., pedestrians), followed by the LVM (e.g., signals on the instrument panel), and lastly the UVM (objects in the distance/the sky). The observed preview asymmetries show that when drivers make saccadic eye movements to different locations, integrating information across these saccades results in more homogenous performance for locations around the visual field.

This study highlights the need in future work to study pre-, post-, and trans-saccadic perception for different saccade directions. Most studies investigating perceptual modulations around saccades, such as pre-saccadic attention (53, 55–57, 59, 69), predictive remapping (70–72), and trans-saccadic integration (3, 7, 9, 29, 63) are based on horizontal eye movements. The preview asymmetries observed here, in line with polar angle differences in pre-saccadic attention (44, 45) call into question the generalizability of these findings from one saccade angle to another.

In summary, the extrafoveal preview effect, as a measure of trans-saccadic integration, follows a reverse pattern of peripheral polar angle asymmetries during fixation. This finding suggests that the visual system actively compensates for polar angle asymmetries in peripheral visual perception by selectively assigning higher weights to the less-well perceived preview information around the time of a saccade. Future neurophysiological studies should investigate how and where such compensation is implemented in the human brain.

## Materials and Methods

### Participants

We recruited 15 participants at NYU-NY. All participants gave informed consent before the experiment, received allowance, and had normal or corrected-to-normal vision. All but two participants (authors XL and NH) were naïve to the purpose of the experiment. One participant was excluded after data collection for having a threshold estimate three SD away from the group average in the valid preview condition (see Procedure). The remaining 14 participants (9 females; all right-handed) had an average age of 28.14 (± 4.74) (M ± SD) years. The experiment was approved by the NYU Abu Dhabi ethical committee and the NYU-NY ethical committee and all experimental procedures were in agreement with the Declaration of Helsinki.

### Apparatus

The experiment was coded with MATLAB PsychToolbox-3 (Brainard 1997), Eyelink toolbox (Cornelissen et al., 2002), as well as the Palamedes toolbox (Prins & Kingdom, 2018) for the PEST staircase procedure (Pentland, 1980), and was run on an Apple iMac Intel Core 2 Duo computer (Cupertino, CA, USA). Stimuli were displayed on a gamma-linearized 20-inch ViewSonic G220fb CRT screen (Brea, CA, USA) (1280×960 screen resolution; 100 Hz refresh rate) with a gray background (luminance: 128 cd/m2). Participants sat at a viewing distance of 57 cm with their head stabilized on a chin-forehead rest. Eye position of the dominant eye was recorded by an EyeLink 1000 Plus eye tracker (SR Research, Ontario, Canada) with the desktop mount, at a sampling rate of 1000 Hz.

### Procedure

The main study included three experimental tasks performed in separate experimental sessions with overall four viewing conditions: Participants first completed a *baseline* task, in which peripheral stimuli were viewed during fixation (**Fig. 1a**), followed by an *integration* task (with viewing conditions *valid* and *invalid* preview intermixed) and a *postview* task. In all tasks, participants judged the orientation of a Gabor patch (clockwise/counterclockwise relative to vertical, 2-alternative forced choice task). To measure contrast sensitivity, stimulus contrast per experimental condition (4 locations × 4 viewing conditions: fixation, valid preview, invalid preview, postview) was separately adjusted by an adaptive staircase procedure (PEST; see SI) based on the correctness of participants’ tilt direction responses.

In all tasks, a white central fixation dot (0.3 dva diameter; dva) and four black placeholders centered at four isoeccentric locations (8 dva) along the upper, lower, left, and right meridian of the fixation dot were presented throughout the trial. Each placeholder consisted of four corners (0.3 dva in length, 0.1 dva in width), indicating the possible locations of upcoming stimuli and potential saccade targets. Each trial started with participants fixating on the central fixation dot. Once we detected continuous fixation (within a radius of 1.5 dva) for 700 ms, the trial continued depending on the specific task.

#### Baseline fixation task

After the initial fixation, four Gabor patches (spatial frequency = 4 cpd; diameter = 2.5 dva; σ = 0.625 dva) were presented at the center of the placeholders for 100 ms while participants kept fixating on the central fixation dot. Three Gabors were vertically oriented distractors, one Gabor was a target tilted ±15° relative to vertical. The location of the target was randomly selected and unbeknownst to the participants until its presentation. Four hundred ms after Gabor offset, the fixation dot color changed to grey and the placeholder at the target location turned white, prompting participants to indicate the tilt direction of the target by pressing a button on a standard keyboard, without time pressure. Audio feedback was provided for 200 ms immediately following the response.

#### Integration task

After the initial fixation, participants (while still fixating at the central fixation dot) peripherally viewed four Gabors, which were presented within the placeholders and independently tilted slightly clockwise or counterclockwise relative to vertical (**Fig. 1b**). The tilt angles of the Gabors were determined by a titration task beforehand (see SI). After 400 ms of preview, a saccade cue (red line of 0.45 dva in length, 0.05 dva in width) superimposed on the fixation dot, indicated to which Gabor participants had to make an immediate saccade. We monitored gaze positions online. As soon as saccade onset was detected (i.e., when two consecutive gaze samples ∼1 ms apart deviated more than 0.18 dva), a visual mask was displayed at each location for 10 ms. This mask (bandpass filtered random noise that covered all possible orientations and with spatial frequency from 1-16 cpd with 38 log-spaced intervals, same contrast and size as the Gabor patch) served to equalize the visual input changes in the valid and invalid conditions (**Fig. 1c**). Note that participants reported to be unaware of the mask due to saccadic suppression (73). The offline saccade detection algorithm used to more accurately detect saccadic eye movements (see Data Analysis section) shows an average interval between actual saccade onset and mask onset of 15.23 (± 5.52) ms (**Fig. 1d**).

Immediately after the mask, the target stimulus was presented at the saccade target. The target stimulus disappeared 80 ms after we detected participants’ gaze within 2 dva from the saccade target center (i.e., the online approximation of a saccade offset). On 98.52% trials, the saccade landed after target stimulus onset, with an average target-saccade interval of 21.42 (± 11.22) ms (**Fig. 1e**). For the remaining <1.5% of trials, target stimulus onset occurred within 20 ms after saccade offset, i.e., within the time range of saccadic suppression (73). After target stimulus disappearance, participants indicated the tilt orientation of the target stimulus. They were explicitly instructed to base their judgment on the post-saccadic target stimulus, which they foveated after the saccade. To prevent participants from impulsively responding to the preview, participants completed at least 36 practice trials with prolonged post-saccadic target duration (300 ms) followed by another 36 practice trials with 80 ms post-saccadic target duration before the *integration* task. We would only start the main task after participants reached 80% correct orientation discrimination performance in the invalid condition of the first practice.

#### Postview task

To measure performance in a neutral, no-preview condition, in this task no Gabors were presented for preview. Participants merely fixated on the central fixation until they saw a saccade cue (the same overall fixation duration as in the *integration* task). Once a saccade was detected, we presented bandpass filtered noise at the target location for 10 ms, followed by a target Gabor at the saccade target. Again, the target Gabor disappeared 80 ms after we detected participants’ gaze within 2 dva from the saccade target center. The rest of the task was identical to the *integration* task.

The study comprised 3 experimental sessions completed over 3-4 days; each lasted 90 minutes. In the first session, participants completed the *fixation* task followed by the first part of the *integration* task. The second session consisted of the *postview* task and the second part of the *integration* task (order counterbalanced among participants). In the third session, participants completed the last part of the *integration* task. The first session of the *integration* task was preceded by a titration task (see SI) to determine the tilt angle of the Gabors used in the *integration* and *postview* tasks. In total, participants performed 2592 trials: 432 trials in both the *fixation* and *postview* tasks, and 1728 trials in the *integration* task. Trials were aborted and repeated at the end of the session if participants 1) did not maintain central fixation during the preview, 2) made a saccade too early (< 100 ms) or too late (> 550 ms) after cue onset, or 3) looked toward a wrong stimulus in an eye-movement task.

### Eye data preprocessing

We analyzed the recorded eye position data and detected saccades (> 2 dva) based on their velocity distribution (74) using a moving average over 20 subsequent gaze samples. Saccade onset and offset were detected when the velocity exceeded or fell below the median of the moving average by three standard deviations for at least 20 ms. Consecutive saccades (onset time within 20 ms) were merged into one saccade. For the analysis of post-saccadic eye movements, we considered eye movements in which the velocity exceeded or fell below the median of the moving average by three standard deviations for at least 2 ms and merged movements with on- and offset difference < 2 ms.

### Data analysis

Contrast sensitivity and saccade parameters were analyzed with R and MATLAB. Contrast thresholds and staircase exclusion were determined with a similar method as in (42; see SI). On average, 12% of staircases were excluded for each participant. Because contrast sensitivity was computed as the reciprocal of the average threshold estimate, it is logarithmically scaled. Therefore, the difference in logarithmic contrast sensitivity between the valid and invalid preview conditions was used as an index for the preview effect. Note that all the findings are the same when the preview effect is defined by the normalized difference (i.e., the difference in contrast sensitivity between the valid and invalid preview conditions divided by their average).

All analyses were performed in R (package ez, version 4.4.0). For all repeated-measures ANOVAs in which the sphericity assumption was not met, we report Greenhouse-Geisser corrected *p-*values. Post-hoc tests for any significant effects were run with the package *emmeans* (version 1.7.4-1 in R). Multiple comparisons were corrected by controlling the false discovery rate. For effect size, we report Cohen’s d for t-tests, *η*_*p*_^2^ for ANOVAs, along with 95% confidence interval (CI) for variance explained by each effect (CI_*d*,_ CI_*η*_). For post-hoc tests and the correlation analysis, we report the 95% CIs for difference and correlation coefficient. We also provide Bayes factors (package *BayesFactor*, version 0.9.12-4.4) for the main findings (i.e., peripheral polar angle asymmetries and preview asymmetries) as a measure of graded evidence.

### Power analysis

We determined the sample size based on previous studies on polar angle asymmetries (e.g., 18, 42, 43, 72) or on the preview effect (e.g., 8, 9, 55). A post-hoc power analysis confirmed that our sample size (n = 14) is associated with 100% power to detect an existing HVA and VMA, and 98% power to detect the preview effect (Cohen’s *d* = 1.18, α = .005), and thus adequate to provide reliable results.

## Acknowledgments

This research was supported by a Marie Skłodowska-Curie individual fellowship (MSCA-IF 898520) by the European Commission to NMH, a National Institutes of Health – National Eye Institute grant R01 EY019693 to MC, and by the NYUAD Center for Brain and Health, funded by Tamkeen under NYU Abu Dhabi Research Institute grant CG012.

## References

1. D. Melcher, C. L. Colby, Trans-saccadic perception. Trends Cogn. Sci. 12, 466–473 (2008).

2. K. Rayner, G. W. McConkie, D. Zola, Integrating information across eye movements. Cognit. Psychol. 12, 206–226 (1980).

3. C. Huber-Huber, A. Buonocore, D. Melcher, The extrafoveal preview paradigm as a measure of predictive, active sampling in visual perception. J. Vis. 21, 12–12 (2021).

4. C. Cont, E. Zimmermann, The Motor Representation of Sensory Experience. Curr. Biol. CB 31, 1029-1036.e2 (2021).

5. E. Zimmermann, R. Weidner, R. O. Abdollahi, G. R. Fink, Spatiotopic Adaptation in Visual Areas. J. Neurosci. 36, 9526–9534 (2016).

6. E. R. Schotter, B. Angele, K. Rayner, Parafoveal processing in reading. Atten. Percept. Psychophys. 74, 5–35 (2012).

7. C. Huber-Huber, A. Buonocore, O. Dimigen, C. Hickey, D. Melcher, The peripheral preview effect with faces: Combined EEG and eye-tracking suggests multiple stages of trans-saccadic predictive and non-predictive processing. NeuroImage 200, 344–362 (2019).

8. A. Buonocore, O. Dimigen, D. Melcher, Post-Saccadic Face Processing Is Modulated by Pre-Saccadic Preview: Evidence from Fixation-Related Potentials. J. Neurosci. 40, 2305–2313 (2020).

9. X. Liu, C. Huber-Huber, D. Melcher, The Trans-Saccadic Extrafoveal Preview Effect is Modulated by Object Visibility in 2022 Symposium on Eye Tracking Research and Applications, (ACM, 2022), pp. 1–7.

10. G. Edwards, R. VanRullen, P. Cavanagh, Decoding Trans-Saccadic Memory. J. Neurosci. 38, 1114–1123 (2018).

11. J. Abrams, A. Nizam, M. Carrasco, Isoeccentric locations are not equivalent: The extent of the vertical meridian asymmetry. Vision Res. 52, 70–78 (2012).

12. M. Carrasco, C. P. Talgar, E. L. Cameron, Characterizing visual performance fields: effects of transient covert attention, spatial frequency, eccentricity, task and set size. Spat. Vis. 15, 61–75 (2001).

13. J. A. Greenwood, M. Szinte, B. Sayim, P. Cavanagh, Variations in crowding, saccadic precision, and spatial localization reveal the shared topology of spatial vision. Proc. Natl. Acad. Sci. 114, E3573–E3582 (2017).

14. M. M. Himmelberg, J. Winawer, M. Carrasco, Stimulus-dependent contrast sensitivity asymmetries around the visual field. J. Vis. 20, 1–19 (2020).

15. M. Jigo, D. Tavdy, M. M. Himmelberg, M. Carrasco, Cortical magnification eliminates differences in contrast sensitivity across but not around the visual field. eLife 12, e84205 (2023).

16. A. S. Baldwin, T. S. Meese, D. H. Baker, The attenuation surface for contrast sensitivity has the form of a witch’s hat within the central visual field. J. Vis. 12, 23 (2012).

17. J. P. Rijsdijk, J. N. Kroon, G. J. van der Wildt, Contrast sensitivity as a function of position on the retina. Vision Res. 20, 235–241 (1980).

18. A. Barbot, S. Xue, M. Carrasco, Asymmetries in visual acuity around the visual field. J. Vis. 21, 1–23 (2021).

19. Y. Kwak, N. M. Hanning, M. Carrasco, Presaccadic attention sharpens visual acuity. Sci. Rep. 13, 2981 (2023).

20. L. Montaser-Kouhsari, M. Carrasco, Perceptual asymmetries are preserved in short-term memory tasks. Atten. Percept. Psychophys. 71, 1782–1792 (2009).

21. C. P. Talgar, M. Carrasco, Vertical meridian asymmetry in spatial resolution: Visual and attentional factors. Psychon. Bull. Rev. 9, 714–722 (2002).

22. J. W. Kurzawski, et al., The Bouma law accounts for crowding in 50 observers. J. Vis. 23, 6 (2023).

23. M. M. Himmelberg, J. Winawer, M. Carrasco, Polar angle asymmetries in visual perception and neural architecture. Trends Neurosci. 46, 445–458 (2023).

24. M. O. Ernst, M. S. Banks, Humans integrate visual and haptic information in a statistically optimal fashion. Nature 415, 429–433 (2002).

25. M. O. Ernst, H. H. Bülthoff, Merging the senses into a robust percept. Trends Cogn. Sci. 8, 162–169 (2004).

26. K. P. Körding, D. M. Wolpert, Bayesian integration in sensorimotor learning. Nature 427, 244–247 (2004).

27. B. Angele, K. Rayner, Parafoveal Processing of Word n + 2 During Reading: Do the Preceding Words Matter? J. Exp. Psychol. Hum. Percept. Perform. 37, 1210–1220 (2011).

28. S. A. McDonald, Parafoveal preview benefit in reading is not cumulative across multiple saccades. Vision Res. 45, 1829–1834 (2005).

29. C. Wolf, A. C. Schütz, Trans-saccadic integration of peripheral and foveal feature information is close to optimal. J. Vis. 15, 1–1 (2015).

30. F. Hutzler, S. Schuster, C. Marx, S. Hawelka, An investigation of parafoveal masks with the incremental boundary paradigm. PLOS ONE 14, e0203013 (2019).

31. M. Carrasco, Visual attention: The past 25 years. Vision Res. 51, 1484–1525 (2011).

32. P. Sharp, D. Melcher, C. Hickey, Endogenous attention modulates the temporal window of integration. Atten. Percept. Psychophys. 80, 1214–1228 (2018).

33. A. Barbot, M. Carrasco, Attention Modifies Spatial Resolution According to Task Demands. Psychol. Sci. 28, 285–296 (2017).

34. A. Fernández, R. N. Denison, M. Carrasco, Temporal attention improves perception similarly at foveal and parafoveal locations. J. Vis. 19, 12 (2019).

35. M. Roberts, B. K. Ashinoff, F. X. Castellanos, M. Carrasco, When attention is intact in adults with ADHD. Psychon. Bull. Rev. 25, 1423–1434 (2018).

36. S. Purokayastha, M. Roberts, M. Carrasco, Voluntary attention improves performance similarly around the visual field. Atten. Percept. Psychophys. 83, 2784–2794 (2021).

37. H. Deubel, The time course of presaccadic attention shifts. Psychol. Res. 72, 630–640 (2008).

38. H. Deubel, W. X. Schneider, Saccade target selection and object recognition: Evidence for a common attentional mechanism. Vision Res. 36, 1827–1837 (1996).

39. N. M. Hanning, H. Deubel, M. Szinte, Sensitivity measures of visuospatial attention. J. Vis. 19, 17 (2019).

40. E. Kowler, E. Anderson, B. Dosher, E. Blaser, The role of attention in the programming of saccades. Vision Res. 35, 1897–1916 (1995).

41. M. Rolfs, M. Carrasco, Rapid Simultaneous Enhancement of Visual Sensitivity and Perceived Contrast during Saccade Preparation. J. Neurosci. 32, 13744–13752a (2012).

42. A. Montagnini, E. Castet, Spatiotemporal dynamics of visual attention during saccade preparation: Independence and coupling between attention and movement planning. J. Vis. 7, 8 (2007).

43. W. J. Harrison, J. B. Mattingley, R. W. Remington, Eye Movement Targets Are Released from Visual Crowding. J. Neurosci. 33, 2927–2933 (2013).

44. N. M. Hanning, M. M. Himmelberg, M. Carrasco, Presaccadic attention enhances contrast sensitivity, but not at the upper vertical meridian. iScience 25, 103851 (2022).

45. N. M. Hanning, M. M. Himmelberg, M. Carrasco, Presaccadic Attention Depends on Eye Movement Direction and Is Related to V1 Cortical Magnification. J. Neurosci. Off. J. Soc. Neurosci. 44, e1023232023 (2024).

46. V. C. Caruso, D. S. Pages, M. A. Sommer, J. M. Groh, Beyond the labeled line: variation in visual reference frames from intraparietal cortex to frontal eye fields and the superior colliculus. J. Neurophysiol. 119, 1411–1421 (2018).

47. J. D. Schall, D. P. Hanes, Neural basis of saccade target selection in frontal eye field during visual search. Nature 366, 467–469 (1993).

48. J. Z. Bakdash, L. R. Marusich, Repeated Measures Correlation. Front. Psychol. 8 (2017).

49. A. Tzelepi, N. Laskaris, A. Amditis, Z. Kapoula, Cortical activity preceding vertical saccades: A MEG study. Brain Res. 1321, 105–116 (2010).

50. M. Rucci, J. D. Victor, The unsteady eye: an information-processing stage, not a bug. Trends Neurosci. 38, 195–206 (2015).

51. M. Nyström, I. Hooge, K. Holmqvist, Post-saccadic oscillations in eye movement data recorded with pupil-based eye trackers reflect motion of the pupil inside the iris. Vision Res. 92, 59–66 (2013).

52. H. H. Li, J. Pan, M. Carrasco, Different computations underlie overt presaccadic and covert spatial attention. Nat. Hum. Behav. 5, 1418–1431 (2021).

53. H. H. Li, A. Barbot, M. Carrasco, Saccade Preparation Reshapes Sensory Tuning. Curr. Biol. 26, 1564–1570 (2016).

54. H.-H. Li, N. M. Hanning, M. Carrasco, To look or not to look: dissociating presaccadic and covert spatial attention. Trends Neurosci. 44, 669–686 (2021).

55. S. Ohl, C. Kuper, M. Rolfs, Selective enhancement of orientation tuning before saccades. J. Vis. 17, 2 (2017).

56. H.-H. Li, J. Pan, M. Carrasco, Presaccadic attention improves or impairs performance by enhancing sensitivity to higher spatial frequencies. Sci. Rep. 9, 2659 (2019).

57. N. De Pisapia, L. Kaunitz, D. Melcher, Backward Masking and Unmasking Across Saccadic Eye Movements. Curr. Biol. 20, 613–617 (2010).

58. J. H. Fabius, A. Fracasso, D. J. Acunzo, S. V. D. Stigchel, D. Melcher, Low-Level Visual Information Is Maintained across Saccades, Allowing for a Postsaccadic Handoff between Visual Areas. (2020). 10.1523/JNEUROSCI.1169-20.2020.

59. D. Melcher, Predictive remapping of visual features precedes saccadic eye movements. Nat. Neurosci. 10, 903–907 (2007).

60. E. E. M. Stewart, M. Valsecchi, A. C. Schütz, A review of interactions between peripheral and foveal vision. J. Vis. 20, 1–35 (2020).

61. S. Kwon, M. Rolfs, J. F. Mitchell, Presaccadic motion integration drives a predictive postsaccadic following response. J. Vis. 19, 12 (2019).

62. M. P. Baumann, A. R. Bogadhi, A. F. Denninger, Z. M. Hafed, Sensory tuning in neuronal movement commands. Proc. Natl. Acad. Sci. 120, e2305759120 (2023).

63. E. Ganmor, M. S. Landy, E. P. Simoncelli, Near-optimal integration of orientation information across saccades. J. Vis. 15 (2015).

64. S. Neupane, D. Guitton, C. C. Pack, Perisaccadic remapping: What? How? Why? Rev. Neurosci. 31, 505–520 (2020).

65. X. Fan, L. Wang, H. Shao, D. Kersten, S. He, Temporally flexible feedback signal to foveal cortex for peripheral object recognition. Proc. Natl. Acad. Sci. 113, 11627–11632 (2016).

66. L. M. Kroell, M. Rolfs, Foveal vision anticipates defining features of eye movement targets. eLife 11, e78106 (2022).

67. O. Jensen, Y. Pan, S. Frisson, L. Wang, An oscillatory pipelining mechanism supporting previewing during visual exploration and reading The temporal constraints during visual exploration and reading. (2021). 10.1016/j.tics.2021.08.008.

68. C. Osterbrink, A. Herwig, Prediction of complex stimuli across saccades. J. Vis. 21, 1–15 (2021).

69. A. Buonocore, A. Fracasso, D. Melcher, Pre-saccadic perception: Separate time courses for enhancement and spatial pooling at the saccade target. PloS One 12, e0178902 (2017).

70. D. Jonikaitis, M. Szinte, M. Rolfs, P. Cavanagh, Allocation of attention across saccades. J Neurophysiol 109, 1425–1434 (2013).

71. X. Wang, et al., Perisaccadic Receptive Field Expansion in the Lateral Intraparietal Area. Neuron 90, 400–409 (2016).

72. J. R. Duhamel, C. L. Colby, M. E. Goldberg, The updating of the representation of visual space in parietal cortex by intended eye movements. Science 255, 90–92 (1992).

73. I. Vallines, M. W. Greenlee, Saccadic Suppression of Retinotopically Localized Blood Oxygen Level-Dependent Responses in Human Primary Visual Area V1. J. Neurosci. 26, 5965–5969 (2006).

74. R. Engbert, K. Mergenthaler, Microsaccades are triggered by low retinal image slip. Proc. Natl. Acad. Sci. 103, 7192–7197 (2006).

75. K. Anton-Erxleben, K. Herrmann, M. Carrasco, Independent effects of adaptation and attention on perceived speed. Psychol. Sci. 24, 150–159 (2013).

