## Supporting Information for "Pre-saccadic Preview Shapes Post-Saccadic Processing More Where Perception is Poor"

### Methods

**Contrast titration procedure.** For each task, Gabor contrast was adjusted on a trial-by-trial basis to determine the Gabor contrasts required for 80% discrimination accuracy. We used an adaptive psychometric staircase procedure (PEST; using the Palamedes toolbox (1) and custom code (https://github.com/michaeljigo/palamedes_wrapper)). In total, there were 16 experimental conditions: 4 viewing conditions (fixation, valid preview, invalid preview, postview) x 4 locations (left, right, upper, lower), for each of which we ran three independent staircase procedures (36 trials per staircase) for each experimental condition in the *fixation* and *postview* tasks, and six staircase procedures for the two viewing conditions in the *integration* task (because we expected a larger within-participant variance). Participants completed the *integration* task in 3 sessions conducted on different days, with each session containing 2 staircases per experimental condition. To achieve a more stable contrast estimate, we “calibrated” the staircases by presenting stimuli at predetermined contrasts for the first 10 trials of each procedure (log-spaced from 1.60% to 63.10%).

The median of the contrast in the last five trials of each individual staircase was used as the threshold estimate. Outliers were defined as threshold estimates deviating more than 0.5 log-contrast units from the average of the other estimates in the same experimental condition of the same individual and were removed. On average, 12% of staircases (1.79% for the fixation task, 8.63% and 20.83% for the valid and invalid conditions in the integration task, 16.67% for the postview task) were excluded for each subject.

**Tilt titration task and results*.*** For the *integration* and *postview tasks*, we adjusted each participant’s tilt angles (relative to vertical) in a separate task using adaptive staircase procedures. The goal was to find the optimal tilt angle for each observer and meridian, which was (1) small enough so pre- and post-saccadic visual inputs in the invalid preview condition (when the Gabor angle flipped to opposite direction during the saccade) were not easily discernible, and (2) big enough so participants were able to perform the orientation discrimination task with ~80% accuracy.

Moreover, when the tilt angle is small (e.g., ~1°), people show an advantage in processing peripheral vertical stimuli on the vertical meridian (VM) over the horizontal meridian (HM) due to radial bias (2). The tilt titration task also served to account for this bias (we used smaller tilt angles for targets at the vertical than horizontal meridian, see below).

Each trial started with 700 ms fixation, followed by a Gabor presented for 50ms within one of the cardinal placeholders. Gabor contrast was fixed at a relatively high level (40%) to minimize the influence of known polar angle asymmetries in contrast sensitivity – which we specifically did not aim to cancel out. The tilt angle was adjusted from 0.25◦ to 10◦ based on a PEST staircase procedure targeting 80% orientation discrimination accuracy. We ran three independent staircases (36 trials per staircase) at each cardinal location. For each participant, we averaged all tilt angle estimates to obtain one value per meridian. These individualized tilt angles were then used in the *integration* and *postview* tasks. The average tilt angle was 1.34° ± 0.48° for HM and 1.09° ± 0.19° for VM.

Note that in line with the radial bias (2), we used smaller tilt angles along the vertical than horizontal meridian. Importantly, this slight but consistent difference in tilt angles (0.26° ± 0.12°; $t(13) = 2.16$, $p = .050$), cannot explain the observed preview asymmetries: With increasing tilt angles (as long as they are < 5°), perceptual judgements are increasingly biased toward the pre-saccadic tilt angle as it diverted from the post-saccadic tilt angle (3). Hence, if our results were influenced by the different tilt angles, we should have seen a larger preview effect along the HM, rather than the VM.

**Results**

**Contrast sensitivity after valid and invalid preview.** We performed a repeated- measures ANOVA on the logarithmic contrast sensitivity with preview validity (valid, invalid) and target location (left, right, upper, lower) as within-subjects factors. We found a main effect of preview validity, $F$ (1, 13) = 42.54*,* $p$ < .001, BF > 100, $\eta_p^2$ = .77, 95% $CI_\eta$ = [.42, .86], contrast sensitivity was higher in valid (31.18 ± 1.91) than invalid condition (14.80 ± 1.79). There was also a main effect of location, $F$ (3, 39) = 4.12, $p$ = .012, BF = 27.34, $\eta_p^2$ = .24, 95% $CI_\eta$ = [.01, .40]. as well as a significant interaction between validity and location, $F$ (3, 39) = 4.28, $p$ = .011, BF = 1.08, $\eta_p^2$ = .25, 95% $CI_\eta$ = [.02, .41], indicating that the preview effect differed among locations (**Fig. S1**).

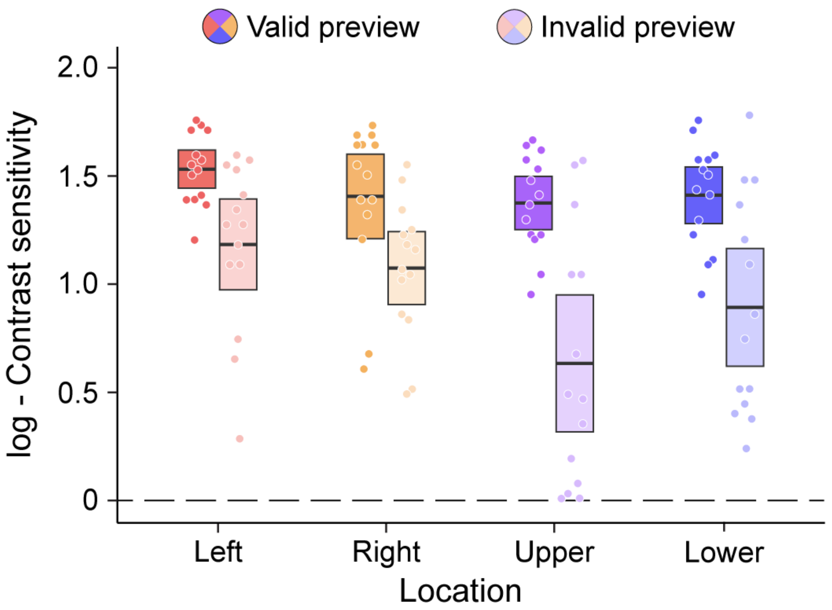


**Fig. S1.** Log-contrast sensitivity (CS) in the integration task as a function or target location, separately for valid (saturated colors) and invalid (transparent colors) trials. CS was computed as the reciprocal of the 80% staircase threshold estimate. Boxes depict the 95% CIs around the mean, represented by the black lines; dots indicate individual observers' log-CS.

### SI References


1. N. Prins, F. A. A. Kingdom, Applying the Model-Comparison Approach to Test Specific Research Hypotheses in Psychophysical Research Using the Palamedes Toolbox. *Front. Psychol.* **9** (2018).
2. Y. Sasaki, *et al.*, The radial bias: a different slant on visual orientation sensitivity in human and nonhuman primates. *Neuron* **51**, 661–670 (2006).
3. E. Ganmor, M. S. Landy, E. P. Simoncelli, Near-optimal integration of orientation information across saccades. *J. Vis.* **15** (2015).
4. R. Engbert, K. Mergenthaler, Microsaccades are triggered by low retinal image slip. *Proc. Natl. Acad. Sci.* **103**, 7192–7197 (2006).
